# Inhibitory-stabilization is sufficient for history-dependent computation in a randomly connected attractor network

**DOI:** 10.1101/2025.10.29.685249

**Authors:** Caelen J. Hilty, Paul Miller

## Abstract

For effective information processing, the response to a sensory stimulus should depend on both the incoming stimulus and the history of prior stimuli. Existing models of neural circuits based on multiple attractor states produced with strong self-excitation can exhibit these properties, but they do not stabilize at biologically realistic firing rates. We demonstrate how a randomly connected inhibition-stabilized attractor network can preserve the computational abilities of recurrent excitatory networks, while stabilizing at arbitrarily low firing rates. Not only does excitatory-inhibitory balance stabilize network activity, inhibitory-stabilization also plays a functional role in history-dependent computation: transient oscillations made possible by inhibitory feedback are sufficient for state-dependent responses to stimulation. Such networks may underlie many cognitive tasks, suggesting a functional role for inhibition-stabilized dynamics in cortical computation.

**Author summary:** General cognitive behavior requires the interpretation of incoming information within its recent context. For example, in sports, a single sensory stimulus: the referee’s whistle, can convey diverse messages: “begin,” “foul,” or “goal” dependent on the events immediately preceding the whistle. We will refer to such situations as “history-dependent,” indicating that the appropriate behavioral response to a given stimulus depends on the prior history of stimulation. History-dependent behaviors include counting, oral communication, and sequence discrimination. In each of these instances, behaviorally relevant information depends less on the characteristics of a single stimulus, but rather on the entire set of stimuli, often including their order. Thus, to perform a wide range of cognitive tasks, the brain must possess a mechanism for short-term memory in which neural responses to a given stimulus depend both on the characteristics of that stimulus and on the recent history of stimulation. Here we study how the dynamics of networks of neurons could support history-dependent behaviors.

## 1 Introduction

General cognitive behavior requires the interpretation of information in context. For example, when processing natural language, the same word can take on different meanings and elicit different responses depending on the words that surround it. Discrimination between sequences of stimuli, stimuli of different durations, or even simple counting requires that the response to a stimulus depend not just on the instantaneous properties of a single stimulus, but on the recent history (or duration) of stimulation. We will refer to such behaviors as history-dependent. To perform history-dependent cognitive tasks, the brain must possess a mechanism for short-term memory in which temporal relationships between stimuli are encoded. Here, we study one candidate mechanism: attractor states in an inhibitory-stabilized network.

Significant experimental evidence suggests that the cortex encodes information in persistent patterns of population-level neural activity termed attractor states (see [1] for review). Hysteretic dynamics, which is highly suggestive of the existence of discrete attractor states in the neural activity landscape has been recorded in multiple model organisms [2, 3]. These neural attractors have been shown to robustly encode task-relevant information. For example, single-unit recordings from prefrontal cortex in macaque display sustained, elevated activity between cue onset and offset only during successful trials on a delayed-response task [4]. Thus, attractor states seem to constitute neural representations of information: attractor states maintain short-term memories.

In response to stimulation or internal noise, neural activity can move between attractor states in a process we name “attractor-state itinerancy” or “itinerancy” for short. Itinerancy is observed experimentally during behavioral trials and is predictive of behavioral responses [5]. In rat gustatory cortex, decision-making about taste palatability progresses through a stereotyped sequence of network states, with sudden transitions between states detectable by single-trial analysis [6, 7]. In a two-alternative forced choice evidence accumulation experiment, cue presentation elicits transitions between states of neural activity in mouse parietal cortex [5]. A sequence of cues causes cortical activity to trace itinerant “walks” through presumed attractor states, culminating in one of many possible final states that predict the animal’s decisions. More specifically, the neural activity demonstrates “state-dependent” itinerancy: the probability of transitions between putative attractor states is modulated by both the identity of the cue and the prior state [5]. Such state-dependent itinerancy is a general mechanism for history-dependent behavior. In responding to a stimulus, the network’s next state is determined both by the properties of an incoming stimulus and the pre-stimulus pattern of network activity. In this manner, the subsequent memory state represents a recent stimulus *given* the information encoded in the prior memory state. Such an encoding of temporal information permits behavioral responses that depend on complex relationships between stimuli like their relative order or number of occurrences.

To study state-dependent itinerancy, we turn to a rate-model of a neuronal circuit, where each variable represents the mean firing activity of a population of neurons. Though this abstraction does discard the intricate response properties of individual cells, it offers analytical and computational tractability and increases model interpretability. The population-level approach of a firing-rate model is an appropriate level of detail to study state-dependent attractor-state itinerancy and the behavior of firing-rate models of inhibition-stabilized networks can be reproduced with populations of model spiking neurons [8, 9].

One challenge of history-dependent computation is that the same signal must both facilitate transitions for some groups of neurons from low activity to higher activity as well as from high activity to lower activity, depending on the pre-stimulus state. Such state dependence requires that the signal can have both an excitatory and an inhibitory impact on a group of cells [10]. In network models composed of self-connected bistable excitatory populations, it was demonstrated that strong short-term synaptic depression—which can alter the relative strength of a stimulus in a history-dependent manner—was sufficient for itinerancy, observed as a switching of activity between multiple states in response to the same stimulus [11–13]. In these models, quasi-stable point attractor states reliably and efficiently represent task-relevant information, including the dynamic characteristics of external stimuli [13]. Stimulus-evoked transitions between attractor states constitute primitive computations, allowing these models to perform evidence accumulation and sequence discrimination tasks with minimal fine-tuning [11]. The depressed synapses of active neurons could allow for the same input that activated them when inactive to also inactivate them when active. However, the high strength of synaptic depression assumed by these models is not well-supported by experimental evidence [14]. Additionally, these models rely on strong recurrent excitatory connectivity to support persistent high-dimensional activity states [11–13]. The dependence on recurrent excitation renders these prior networks vulnerable to runaway activity, such that the most active neurons in a state fire at rates higher than those typically observed [3–5].

To address the biological realism of itinerant multi-attractor networks, here we make use of the experimental evidence suggesting that cortical networks are characterized by similarly strong feedback connections, but recurrent inhibition prevents the recurrent excitation from producing spontaneous runaway activity. Such networks are inhibition-stabilized [15–17] whereby the unstable excitatory subnetwork can be stabilized at arbitrarily low firing rates by the strong inhibitory feedback. A typical feature of inhibition stabilization is a “paradoxical” response to stimulation, where excitation to the inhibitory subnetwork decreases both excitatory and inhibitory activity [15, 17]. The paradoxical response is sufficient evidence for inhibitory stabilization [18] and has been observed in rat hippocampus [17], mouse neocortex [16], ferret visual cortex [19], and cat visual cortex [15], using a variety of perturbation techniques. Additionally, loss of cortical inhibition has been causally related to epileptic seizures in cats [20] and mice [21], which suggests that inhibition stabilization is necessary to constrain runaway excitation in cortex. While inhibition stabilization may not be present in all network states [22], these observations provide ample evidence for its ubiquity in neocortex. We hypothesized that a model of a multiple-attractor network that accounts for these observations could overcome the limitations of prior models and indicate a strong functional role for inhibitory stabilization in cognitive processes.

The canonical mathematical model of inhibitory-stabilization is composed of a single excitatory and single inhibitory population with strong connections within and between populations [15, 17]. These minimal circuits can be tuned for bi-stability, transitioning bidirectionally between a DOWN state and an inhibition-stabilized UP state in both firing rate models [8] and spiking network models [23]. Here we propose a novel point-attractor model composed of many canonical inhibition-stabilized pairs (ISPs) in the bistable regime – an “inhibition-stabilized network” (ISN). We hypothesize that inhibition-stabilized dynamics could provide a biologically plausible negative feedback mechanism to permit state-dependent responses to stimulation in addition to their canonical role in stabilizing network activity at lower, more biologically realistic firing-rates. We show that inhibitory stabilization can result in transient oscillatory dynamics around the UP states. We exploit these dynamics to facilitate state-dependent transitions between quasi-stable attractor states in response to external stimulation. Coupling many ISPs with weak, random cross-connectivity produces a multiplicity of high-dimensional attractor states. With sufficient heterogeneity in the weak cross-connectivity, the ISN can exhibit state-dependent responses to external stimulation, transitioning between these quasi-stable attractor states.

We formally treat our attractor networks as biologically plausible instantiations of finite state automata. In such automata, also known as finite-state machines, inputs cause transitions between a finite set of discrete states such that distinct sequences of repeated inputs may terminate in distinct final attractor states. We demonstrate how random inhibition-stabilized attractor networks can implement state machines that solve a simple two-alternative forced choice evidence accumulation task adapted from [5]. Network performance captured key behavioral features: sigmoidal psychometric curves as well as primacy and recency effects.

Our model describes a hypothesis for how a large class of computational problems, those solvable by finite-state machines, could be solved by a biologically realistic neural system, bridging approaches from neuroscience and the theory of computation.

## 2 Models

An ISN is composed of *N* canonical inhibition-stabilized E-I pairs of rate units (ISPs) and is governed by the equation:

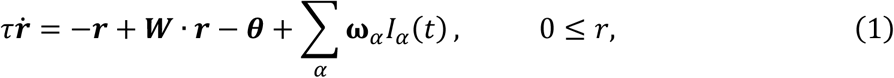

where τ is the time constant for the mean firing rate (10 ms for all simulations), ***r*** is a vector of length 2*N* containing the mean firing-rate of each unit, ***W*** is a 2*N*-by-2*N* connection weight matrix, ***θ*** is a 2N-vector of activation thresholds for each firing-rate unit. 𝐼_𝛼_*(t)* is a scalar of time-dependent input current arising from stimulus 𝛼, with the stimuli distinguished via the weight-vector 𝛚_𝛼_. In situations where we consider only a single stimulus, 𝛚_𝛼_ is the unit vector and identical time-dependent input current is supplied to all units.

Constraints on the weight matrix distinguish the ISN from other possible recurrent neural networks. The weight matrix is parameterized by a mean and standard deviation of weights for each possible type of connection between ISPs (E-to-E, E-to-I, etc.): we exclusively consider networks where each isolated pair is identical. To generate ***W***, we proceed row-by-row, sampling from a log-normal distribution and enforcing Dale’s law. Each row of the matrix is resampled until its statistics exactly match the specified parameters, such that statistics of each row are identical. Inhibition-stabilized dynamics within an ISP are achieved by enforcing structure onto this random matrix: we specify the weights of the comparatively strong connectivity within each pair. The connectivity within each pair is constrained to display the characteristic paradoxical response to stimuli: the inhibitory nullcline must have a higher gain than the excitatory nullcline [8]. The overall effect is that each unit receives the same total weight of incoming connections, which is dominated by the within-pair connections to produce inhibitory stabilization.

While analytic treatment of networks with N > 1 is feasible, we turn to numerical simulations for the bulk of this study. Network activity is approximated by numerically integrating the governing equations with a timestep of 10 nanoseconds using a modified Runge-Kutta (4th Order) method [24] that enforces the threshold nonlinearity after every update step.

All codes used in this paper are available at: https://doi.org/10.5281/zenodo.17470740.

## 3 The single inhibition-stabilized pair

### 3.1 Analytical treatment of a single ISP

In the case where N = 1 (a single ISP), we can explicitly write out the product of the weight matrix and the firing-rate vector and the network reduces to the canonical form:

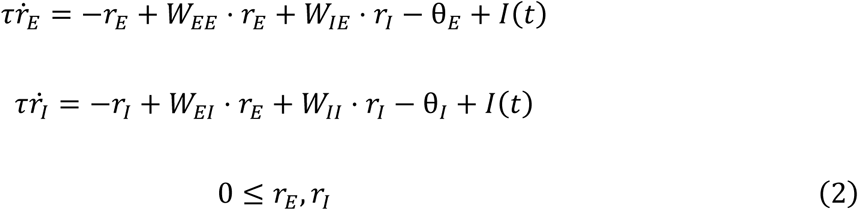

where 𝑊_𝐸𝐸_ corresponds to the connection from the excitatory unit to itself, 𝑊_𝐼𝐸_ is the connection from the inhibitory unit to the excitatory unit, and so on. The non-zero fixed point of equation 2 is given by:

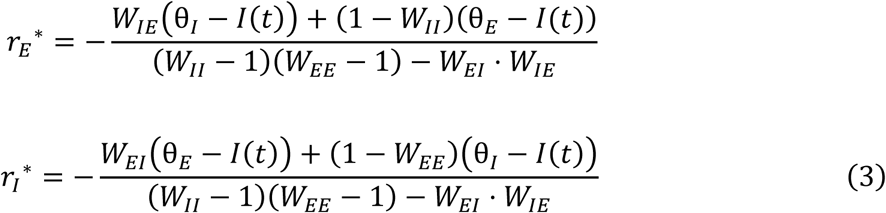

and the origin is a fixed point when θ_I_, θ_E_ ≥ 0 (see Supplemental Materials). The Jacobian of equation (2), evaluated at the non-zero fixed point is:

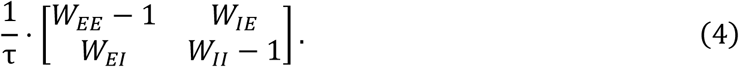

Thus, the trace and determinant of the Jacobian are:

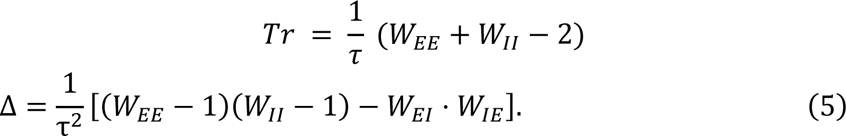

The non-zero fixed point is stable when Tr < 0 and Δ > 0. The fixed point is a stable node when Tr^2^ > 4Δ and a stable spiral when Tr^2^ < 4Δ: the latter will prove to be important for producing itinerancy (Fig. 1 and dashed line in Fig. 2A). The origin is locally asymptotically stable when θ_I_ and θ_E_ are strictly negative: these terms dominate the dynamics when x and y are sufficiently close to zero and thus negative values force trajectories to the origin.

**Figure 1.**
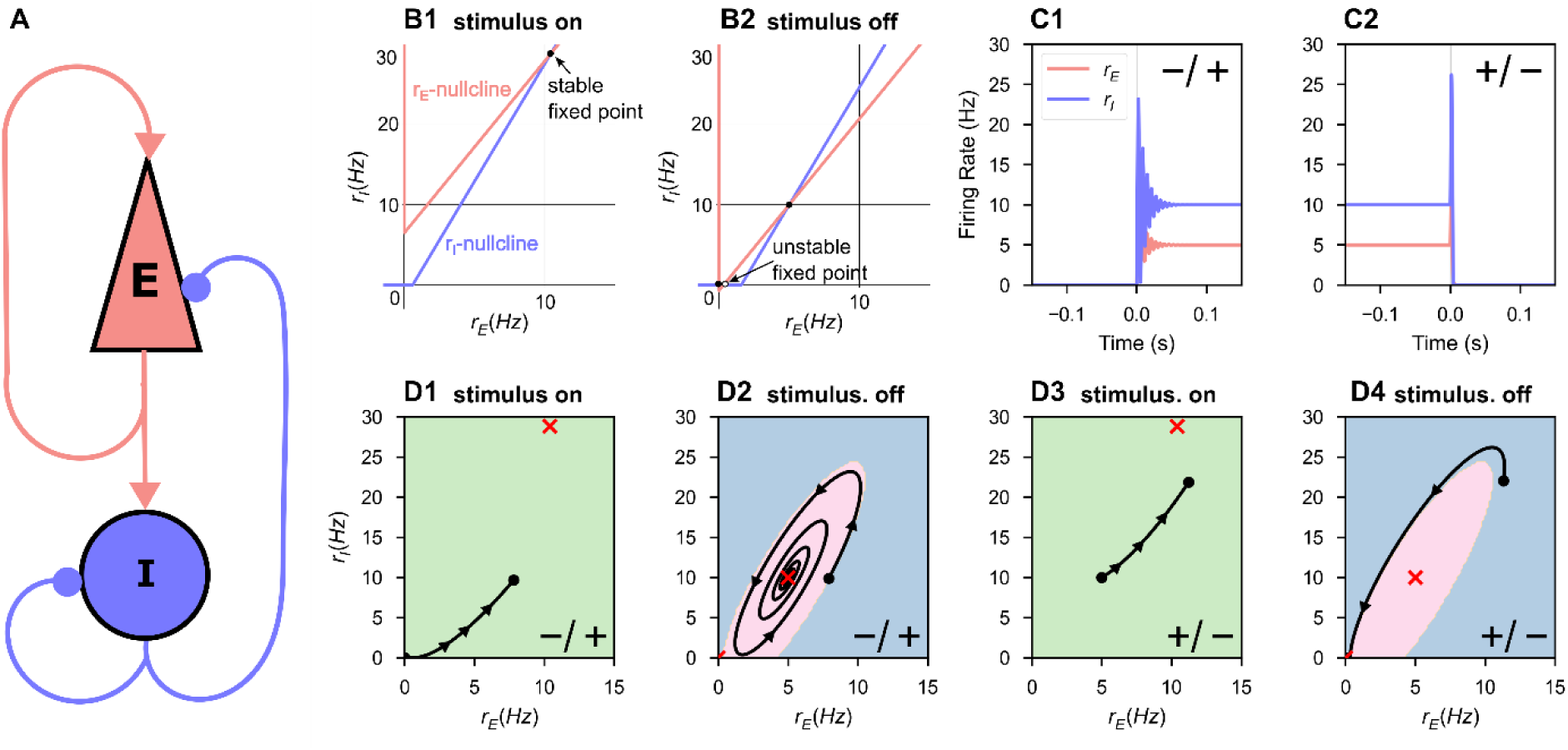
A repeated stimulus elicits bidirectional UP-DOWN state switches in a single bistable ISP. A fixed stimulus (I_app_ = 52.15, τ_dur_ = 1 ms) was repeatedly applied to a single bistable inhibition-stabilized pair. **A.** Schematic of the connected pair with excitatory unit (red) and inhibitory unit (blue). **B.** The nullclines of the system under stimulus (**B1**) and in the absence of stimulus (**B2**). During the stimulus, the ISP is perturbed out of the bistable regime. **C.** Rate-unit responses to stimulus application, time locked to stimulus onset, beginning from the DOWN state (**C1**) and from the UP state (**C2**). **D.** The same trajectories as in **C** plotted in rate-space. Background color indicates the boundaries of the basin(s) of attraction for the attractor state(s), which are indicated by the red cross(es).

**Figure 2.**
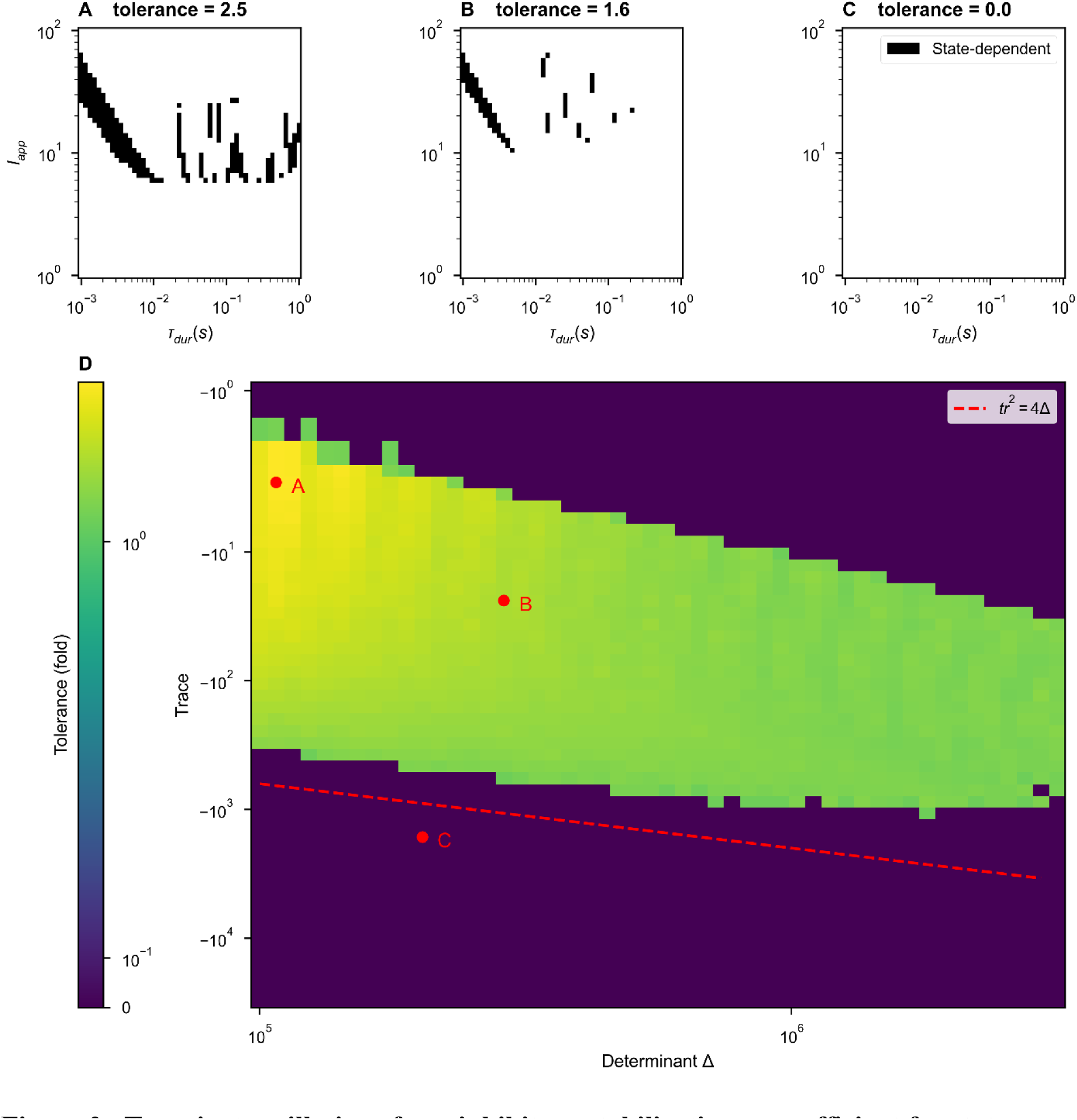
Transient oscillations from inhibitory stabilization are sufficient for state-dependence in a single inhibition-stabilized pair. **(A-C)** For N=1 bistable networks (single ISPs) described by points in **(D)**, we swept over stimulus amplitudes and durations, applying two stimuli. Where the plot is black, stimuli elicited transitions both from DOWN to UP and UP to DOWN *i.e.* state-dependent responses were observed. Example dynamical traces can be found in Supplemental Figure 1. **(D)** The range of stimuli able to produce distinct state-dependent responses for individual bistable pairs as a function of determinant and trace of the Jacobian of the governing equations (Eq. 2). Fixing the firing thresholds (θ_E_ = 5.34, θ_I_ = 82.43) and the firing rates at the stable UP state (*r_E_\** = 5 Hz, *r_I_\** = 10 Hz), we numerically solved for the remaining ISP parameters at each pairwise combination of determinant and trace. With each ISP, we applied a wide range of stimuli and recorded the percent of stimuli over which the network displayed state-dependent responses. Color indicates the stimulus tolerance, which is an approximation in the fold variation in stimulus features that produce state-dependent responses. The dashed red line indicates the boundary between stable fixed points and stable spirals (see Eq. 5 and following text).

The N = 1 network is bistable when 1) the inhibitory threshold is greater than the excitatory threshold and both are greater than zero (θ_I_ > θ_E_ > 0) and 2) the inhibitory gain is greater than the excitatory gain [8] (also see Supplemental Materials). In the threshold-linear model studied here, these conditions produce a stable DOWN state with no firing and a stable UP state with non-zero firing rates in both populations (Fig. 1B.2). In this bistable regime, if the same such stimulus can elicit both a DOWN-to-UP and an UP-to-DOWN state transition, we identify the network’s response to that stimulus as state-dependent.

### 2.2 Inhibition stabilization is sufficient for state-dependence

We studied the response of the N = 1 network to a single stimulus: a rectangular current pulse applied with the same magnitude to both the excitatory and inhibitory population, *i.e.* assuming no structure or cell-type specific segregation of inputs. The current takes the form:

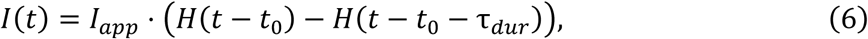

where 𝐼 is the current, 𝐼_𝑎𝑝𝑝_ is the amplitude of the pulse, *H* is the Heaviside function, 𝑡_0_ is the stimulus onset, and τ_𝑑𝑢𝑟_is the duration of the stimulus.

A transient stimulus with magnitude greater than the excitatory threshold is sufficient to temporarily destabilize the DOWN state (Fig. 1B.1, 1D.1), causing a DOWN-to-UP state transition (Fig. 1C.1, 1D.1-2). The same stimulus can elicit an UP-to-DOWN state transition by temporarily moving the UP state to a new location in rate-space (red ‘X’, Fig. 1D), towards which the network activity evolves (Fig. 1D.3). If the stimulus terminates when the network is in the DOWN basin of attraction, an UP-to-DOWN transition occurs (Fig. 1C.2, 1D.3-4).

**Figure 3.**
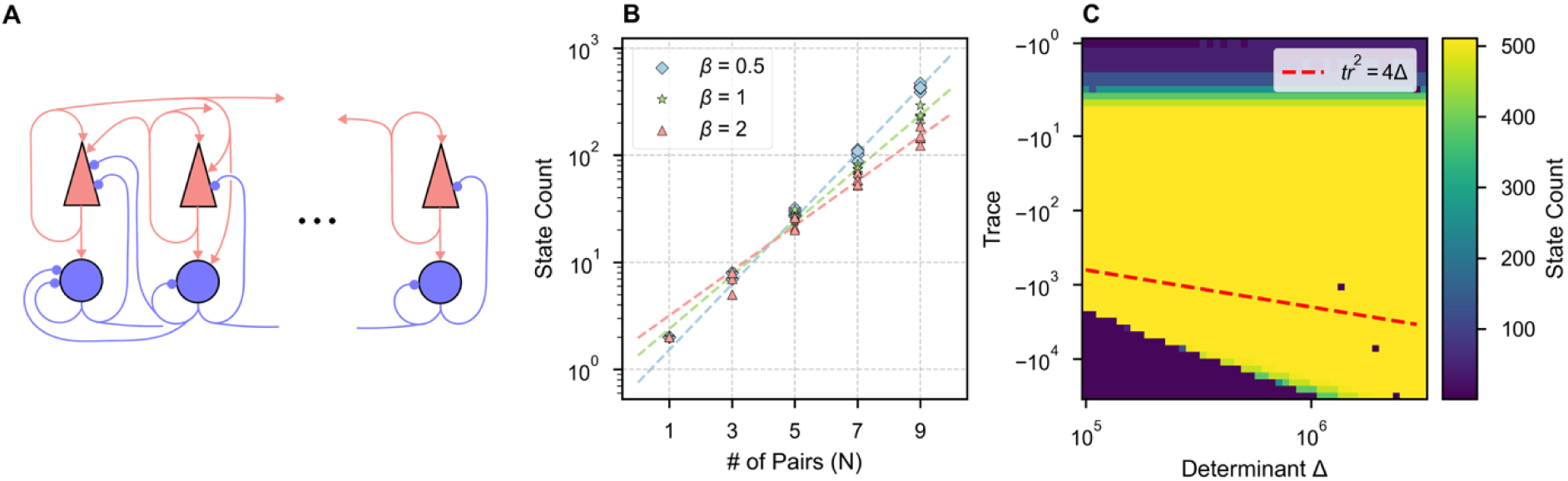
ISNs can exhibit multi-stability robust to variation in within-pair connectivity. **(A)** Schematic of many inhibition-stabilized pairs connected all units to all units. **(B)** Multi-stability grows exponentially with network size. Ten random cross-connectivity matrices were tested at each network size (symbols) for each of three levels of mean cross-connection strength (𝛽 scales the fiducial mean in the Supplemental Materials). The data were fit with an exponential curves (dashed lines), corresponding to 𝑦 = 0.75 × 2.02^𝑥^, 𝑦 = 1.34 × 1.78^𝑥^, 𝑦 = 1.96 × 1.62^𝑥^, for 𝛽 = 0.5, 1, 2 respectively, indicating fewer states with stronger cross-connections. **(C)** The number of attractor states, indicated by color, of an N = 9 network with homogenous cross-connectivity and 𝛽 = 1, as a function of the trace and determinant of the Jacobian at the UP-state for each individual ISP, as obtained by varying within-pair connections with fixed firing rate of UP state. Note the maximum number of states is 𝟐^𝑵^ = 𝟓𝟏𝟐 and is obtained across a wide range of intra-pair connection strengths. Spiral dynamics occur above the red-dashed line.

**Figure 4.**
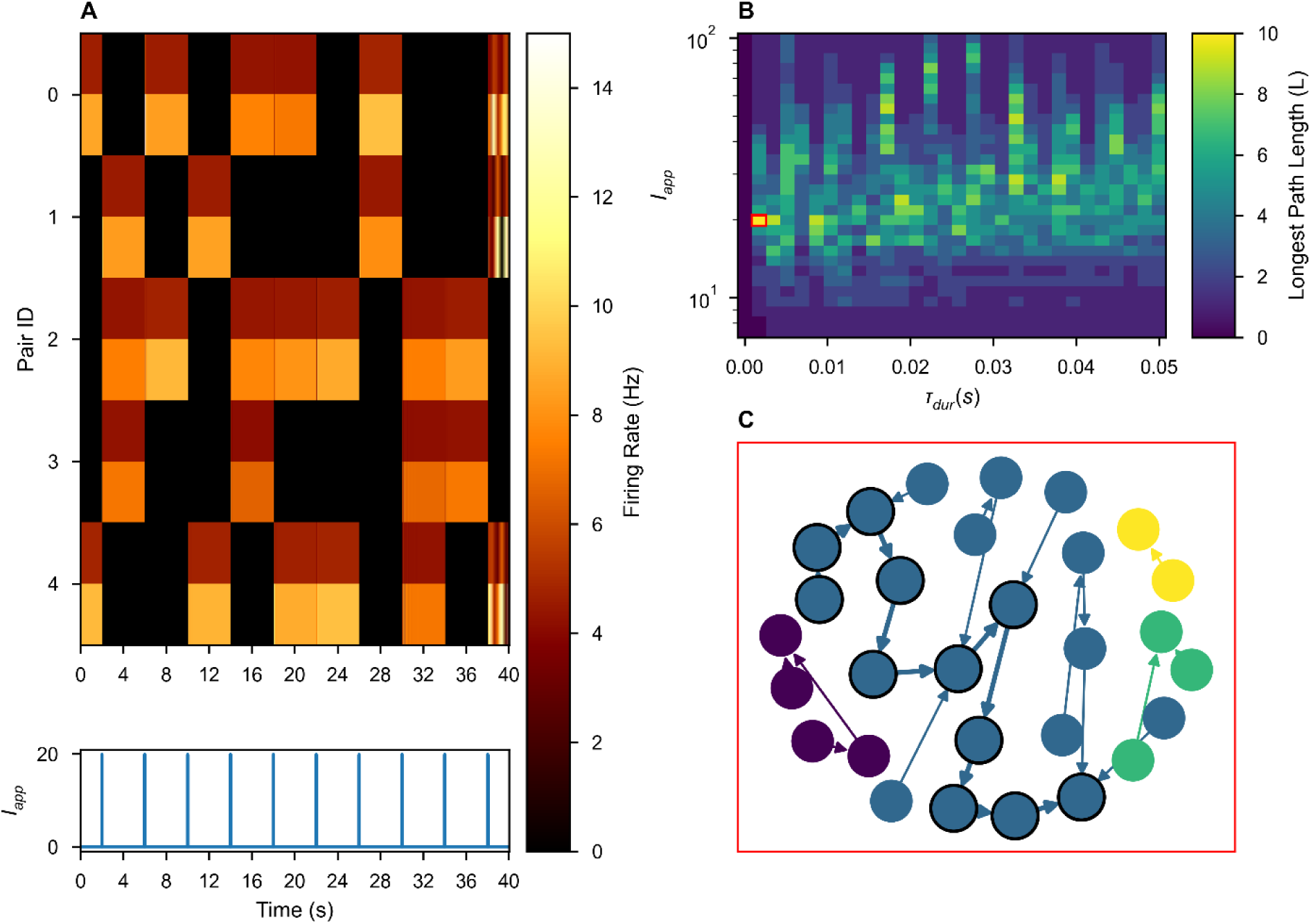
Multi-stable ISNs with weak heterogeneous cross-connections exhibit itinerancy that is robust to variation in stimulus. **(A)** Network activity over time in response to a repeated uniform stimulus with τ_dur_ = 19.9 and I_app_ = 1.7 ms. In each pair the firing rate of the excitatory unit has lower rate and is depicted above that of the inhibitory unit. **(B)** The duration and amplitude of the applied stimulus was varied continuously (log axis for amplitude) and the length (L) of the longest itinerant path elicited was recorded, represented as color. The pair of stimulus parameters used in **(A)** and **(C)** is outlined in red. **(C)** Network itinerancy is visualized in a graph, where each node indicates a network state and each edge indicates a stimulus-evoked transition. Each weakly connected subgraph is identified by a unique color. One of the longest paths (bold edges, black outlined nodes) is shown in **(A)**.

Next, we continuously varied network parameters and evaluated each network’s selectivity of history-dependent responses to the characteristics of the applied stimuli. We varied the stimulus amplitude and duration, applying each stimulus (an amplitude-duration pair) to the network twice, once in the DOWN state and once in the UP state, recording whether the stimulus elicited a state-dependent response – a transition from DOWN to UP and from UP to DOWN. Figure 2A-C summarize the results of such a sweep for three example networks. We observed state-dependent responses for a range of stimulus and network parameters (Figure 2A-B).

To quantify the robustness of network responses, we converted *f_state-dependent_*, the fraction of stimuli for which a state-dependent response was observed, to a scale- and sampling-invariant measure of robustness, “tolerance,” via the following formula:

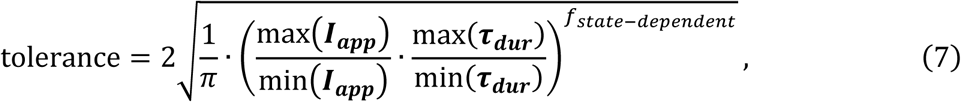

where the boldface stimulus parameters ***I_app_*** and ***τ_dur_*** denote the sets containing the properties of all stimuli simulated for which any state-dependent responses were observed. Tolerance treats the state-dependent region of stimulus space as a continuous circle on log-log axes, reporting the diameter of that circle in units of fold-variation.

While we observed distinct state-dependent responses in many parameter regimes, state-dependence is a non-trivial network property. Even when tested across a thousand-fold range of stimulus durations and a hundred-fold range of stimulus amplitudes, some inhibitory-stabilized, bistable pairs did not display state-dependence, corresponding to a tolerance of zero (Figure 2C).

By inspection of Figure 1, we hypothesized that stable spiral fixed points were necessary for state-dependent responses to stimuli in a bistable ISP. For a stimulus to elicit a DOWN-to-UP transition, that is perturb an entirely quiescent network into activity, the input must be excitatory. But for state-dependence, the same excitatory stimulus must also silence activity in the UP state. Thus, a negative feedback mechanism is necessary for state-dependence [10, 13]. If the dynamics around the UP state are simply nodal, the excitatory input that increases activity from DOWN-to-UP and further increases activity from the UP state is unable to generate a final state of low activity. However, we hypothesized that spiral dynamics could respond to excitation with a transient decrease in activity sufficient to push the network into the DOWN basin of attraction at the time of stimulus termination, as observed in the spiral rotating the activity around the temporary, stimulus-induced non-zero fixed point in Fig. 1C.4.

To test our hypothesis, we continuously varied the character of the UP fixed point from a stable node to a stable spiral, affecting the shapes of trajectories around UP fixed points in both the autonomous and forced system. In Figure 2D, we varied the trace (Tr) and determinant (det) of the Jacobian matrix for Eq. 2, evaluated at the UP state. When 𝑇𝑟 > 4 · 𝑑𝑒𝑡, the UP state is a stable node. When 𝑇𝑟 < 4 · 𝑑𝑒𝑡, the UP state is a stable spiral. We chose fixed values for the firing rates of the UP state (with 𝑟_𝐼_ > 𝑟_𝐸_) and the excitatory and inhibitory thresholds. Together, these constraints (fixed trace, determinant, thresholds, and steady-state firing rates rates) define a system of nonlinear equations over the weights of the within-pair connectivity, which we solved numerically using a root solver from SciPy [25] (see Supplementary Materials). Each solution describes a unique ISP, and – because of our choice of fixed firing thresholds – all solutions shown fall in the bistable inhibition-stabilized regime.

For each trace-determinant pair in Fig. 2D, we found such a solution. For each set of network parameters, we evaluated network tolerance as in Fig. 2A-C. We did not observe distinct state-dependent responses to stimuli in the purely stable regime: the UP state must be a stable spiral for history-dependent responses to stimulation (Fig. 2A). Within the spiral regime, we observed significant stimulus tolerance within a large range of the parameter space sampled. The upper-right region lacking history-dependent responses is an artifact of numerical simulation: the system did not decay to within a 0.1 Hz radius of a fixed point within 12 seconds.

Thus, inhibitory stabilization can produce spiral dynamics that permit an UP-to-DOWN transition in response to excitation. Without transient oscillatory behavior, the model’s threshold-linear dynamics impose no upper bound on the basin of attraction for the UP state (Fig. 1). In prior work, short-term synaptic depression could impose such an upper bound but was unrealistically strong [11–13]. In our work here, no negative feedback dynamics other than inhibitory-stabilization are present in the model: the intrinsic oscillations arising in the inhibitory-stabilized stable-spiral regime can produce distinct state-dependent responses.

## 4 The inhibition-stabilized network exhibits robust itinerancy between multiple attractor states

A simple binary memory is insufficient for encoding information during most cognitive tasks. Here we consider networks with N > 1 ISPs and multiple high-dimensional attractor states (Fig. 3A). In the networks considered below, parameters were selected such that each ISP would be bistable if isolated from the network, while pairs were connected weakly, all-to-all with all four types of excitatory/inhibitory cross-connections. Each attractor state can be uniquely characterized by the identities of the ISPs in their UP states. Therefore, a network of N pairs possesses a maximum of 2^N^ states if every possible permutation of UP/DOWN states for each pair is stable.

Indeed, we observed that increasing the number of ISPs in the network can result in a combinatorial explosion of fixed points (Fig. 3B). We vary the number of pairs in a fiducial ISN, where parameters (defined in the Supplementary Material) are selected such that each pair would be bistable if isolated from the network with stable rates of 𝑟_𝐸_ = 𝑟_𝐼_ = 0 or 𝑟_𝐸_ = 5Hz, 𝑟_𝐼_ = 10Hz. As the network increases in size (N), we scale the mean and standard deviation of the distribution of cross-connections by a factor of 1/N, which keeps the mean total input to each unit constant and maintains a constant coefficient of variation in the distribution of connection strengths (while the total input to each unit remains identical). At each network size, we generate five new random cross-connectivity matrices and record the number of network states. The number of discrete attractors in each network is estimated by sampling 3·2^N^ initial conditions, corresponding to initializing the network with each possible permutation of *k* active units for all *k* ≤ *N* with three different guesses for the stable rates of the UP units. After initialization, we allow the network to stabilize and record the number of unique final states. We find that the relationship between network size and number of states is fit by an exponential curve of the form:

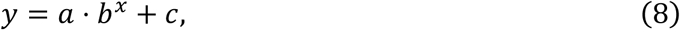

where x is the number of pairs and y is the number of states (Fig. 3B, dashed line).

Multi-stability is robust to variation in the character of the UP state of each individual ISP. Taking for granted the assumption that dynamics are dominated by within-pair connections, we varied the trace and determinant of the Jacobian at the UP state for each ISP only by modifying the within-pair connections, exactly as in Figure 2. At each solution, we performed simulations to count network attractor states (Fig. 3C). Here, we treated a network with the same mean cross-connectivity as in Figure 3B, but with no variability in the cross-connectivity—*i.e.,* the cross-connection strength between all pairs is identical. This reduced the time-complexity of the state-counting algorithm from O(2^N^) to O(N): if there is a stable attractor with n ≤ N pairs in the UP state, then all permutations of n pairs in the UP state are possible because every pair is identical. Thus, we only sampled 3N initial conditions, corresponding to initializing the network with only one possible permutation of *k* active units for all *k* ≤ *N* with three different guesses for the stable rates of the UP units. We observed no dependence of multi-stability on the dynamics around the stable UP state.

The extensive multi-stability possible in some networks endows the model with a large memory capacity. However, memory alone is insufficient for history-dependent cognitive tasks. We evaluate “network itinerancy,” the ability of a network to transition between attractor states in response to external stimulation, in networks with randomness in their cross-connections.

Operationally, a network displays history-dependent itinerancy if, in response to the *same* stimulus, the network transitions between a sequence of many attractor states (Fig. 4A). To evaluate a network’s itinerancy, we initialized it in each of its attractor states, then injected a rectangular pulse of current into all rate-units with uniform stimulus amplitude and duration. A network’s possible responses to a single stimulus can be represented as a graph, where each node is an attractor state and each edge is the state transition elicited by that stimulus. The length of the longest path through such a graph (Fig. 4C) offers a discrete measure of the “magnitude” of network itinerancy, which we measure as a function of the stimulus parameters (Fig. 4B).

Network itinerancy is robust to these perturbations—so long as stimuli are sufficiently strong to cause a change in network activity and not so strong that they dominate all prior activity in the network—circuits display significant itinerancy for a wide range of possible stimuli.

## 5 The inhibition-stabilized network implements a finite-state machine

For a rigorous framework in which to understand how itinerant paths might perform computations in a high-dimensional attractor network, we appeal to the theory of computation. Finite state-dependent information processing algorithms can be solved by a class of automata known as finite-state machines [26]. A deterministic finite state machine (FSM) is defined by: a finite set of states 𝑸, a fixed initial state 𝑞_0_ ∈ 𝑸, a finite set of input symbols (an “alphabet”) 𝚺, a state-transition function (𝑸 × 𝚺) → 𝑸 which describes how inputs result in transitions between states, a finite set of outputs, ***O***, and an output function ***Q →O*** that assigns output labels to each state [26].

Here, we explicitly treat our inhibition-stabilized point-attractor network as a biological finite-state machine. We simulated an N = 5 network’s responses to sequences of similar stimuli during a simple evidence accumulation task (Fig. 5). In experiments by Morcos and Harvey [5], mice ran down a T-shaped track in virtual reality and were presented with six total visual cues on the left and right at fixed intervals (Fig. 5A). At the track intersection, mice were trained to turn towards the direction with more cues. Mice were demonstrated to use evidence from multiple cues to make their decisions, thus displaying history-dependent behavior [5]. To simulate this task, we presented the network with a sequence of six stimuli. We defined two possible stimuli to compose these sequences, representing a left and a right cue. The two stimuli were rectangular pulses of current with the same amplitude and duration, distinguished by their distinct random vectors of weights, which modulated the input current to each unit in a stimulus-dependent manner. Elements of the weight vectors, ***ω_cue,_*** were drawn from a standard log-normal distribution, then rescaled such that each weight vector had a unit mean. These weights represent the inputs from sensory-processing regions of cortex.

**Figure 5.**
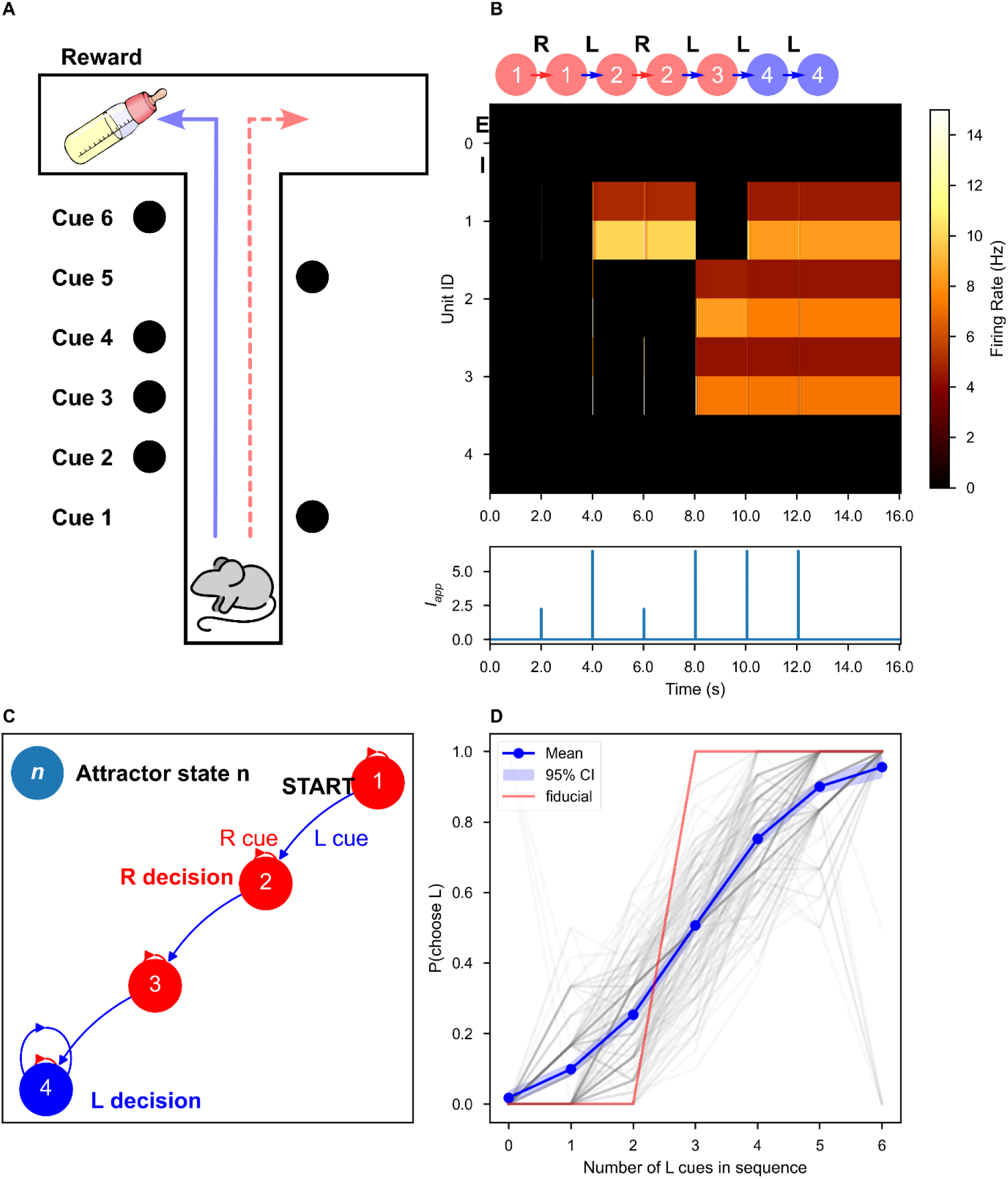
The inhibition-stabilized network implements a finite state machine to solve an evidence-accumulation task. **(A)** Schematic of the behavioral experiment from Morcos and Harvey [5]. As the mouse progresses down the track, visual cues were presented at fixed intervals. Blue line indicates a correct decision (left) for this example. Dashed red line indicates an error. **(B)** An example network response to a sequence of stimuli (*τ_dur_* = 10 ms, *I_app_* = 10.12). Top: Graphical representation of state transitions in response to a sequence of two stimuli, where each node is an attractor state and each edge represents stimulus application. The accepting states are indicated by the color of the node: *e.g.,* a sequence that terminates in a red node results in a right turn decision. Middle: firing rate of each unit over time, pair by pair, encoded as color. Bottom: the external input to the first unit over time. **(C)** The full network response to all possible sequences of stimuli, visualized as a state machine diagram. Directed edges indicate the network response to the left (“L”) or right (“R”) stimuli. The accepting states are indicated by the color of the node. **(D)** Psychometric curves for 1000 random pairs of stimulus weight vectors. For each stimulus pair with reliability above 0.73 (a level achievable without state-dependence by consideration of only a single stimulus), the probability of a left-turn decision given the number of left-turn cues in the sequence was computed. The curves are better fit by a logistic regression than by a linear regression (Supplemental Materials). The red curve indicates the psychometric curve for the example state machine shown in **(B)** and **(C)**

Network responses to sequences of stimuli during this task can be understood as an implementation of an FSM (Fig. 5). Our attractor networks possess a finite number of discrete states. For the fixed initial state of the FSM, we selected the quiescent network state (which is always stable in our formalism). The set **Σ** = {**ω_L_**, **ω_R_**} is contains the possible inputs to the network where **ω_L_** and **ω_R_** represent the left and right cues, respectively. The state-transition function is encoded in the dynamics of the network, so we used a modified breadth-first search algorithm to discover it: for each state, we simulated the response to both stimuli, followed by an equilibration period, and recorded the new stable states (Fig. 5B). If the new states had not yet been traversed, we repeated this process until no further new states were found. The state transition function can be visualized as a graph, where each node is a state and each edge is an element of the stimulus alphabet (Fig. 5C).

The final network state in response to a sequence of left and right stimuli can be determined by tracing the state-machine graph. The ideal network for this task has a unique final response that discriminates between sequences with more left cues and sequences with more right cues.

We assume that an optimal output function **Q →** {“left turn”, “right turn”} could be learned by reinforcement if the corresponding states for the distinct behavioral responses are separated enough (as in [11, 27, 28]). We procedurally found the globally optimal output function to decode the final network states in response to every possible sequence of stimuli. We define a metric of network performance, “reliability,” which measures deviation from the perfect separation of sequences. Consider a deterministic attractor network with *n_states_* attractor states.

Let *S_i_* be the set of sequences for which the network terminates in state *i* ≤ *n_states_*. Let *L* and *R* be the sets of sequences for which the left or right cue is more frequent, respectively. Note that *L* and *R* do not include sequences with an equal number of left and right cues. Then:

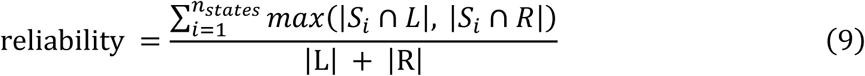

If no sequence discrimination occurs, the network achieves a reliability of 0.50, whereas a perfect network—in which each final state is reached by a subset of sequences either with only more right than left cues or with only more left than right cues—achieves a reliability of 1.00. A network which makes its decision purely based on a single stimulus achieves a reliability of 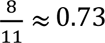 0.73, which we treat as a baseline: any network achieving greater than 0.73 reliability must process sequences in a history-dependent manner.

The example network in Figure 6 implements one of the state-machines that perfectly solves the task with just four states, the minimum required for perfect performance. To demonstrate history-dependent task performance, we varied the task by choosing 1,000 new pairs of random weight vectors to represent the left and right stimuli. For the pairs for which the network achieves above baseline reliability, we measured the probability of a left turn choice as a function of the number of left cues in each sequence, producing a psychometric curve (Fig. 5D). For networks with above-baseline performance, this curve is significantly better fit by a logistic regression than linear regression (see Supplemental Materials, Figure 2), in agreement with the experimental data [5].

**Figure 6.**
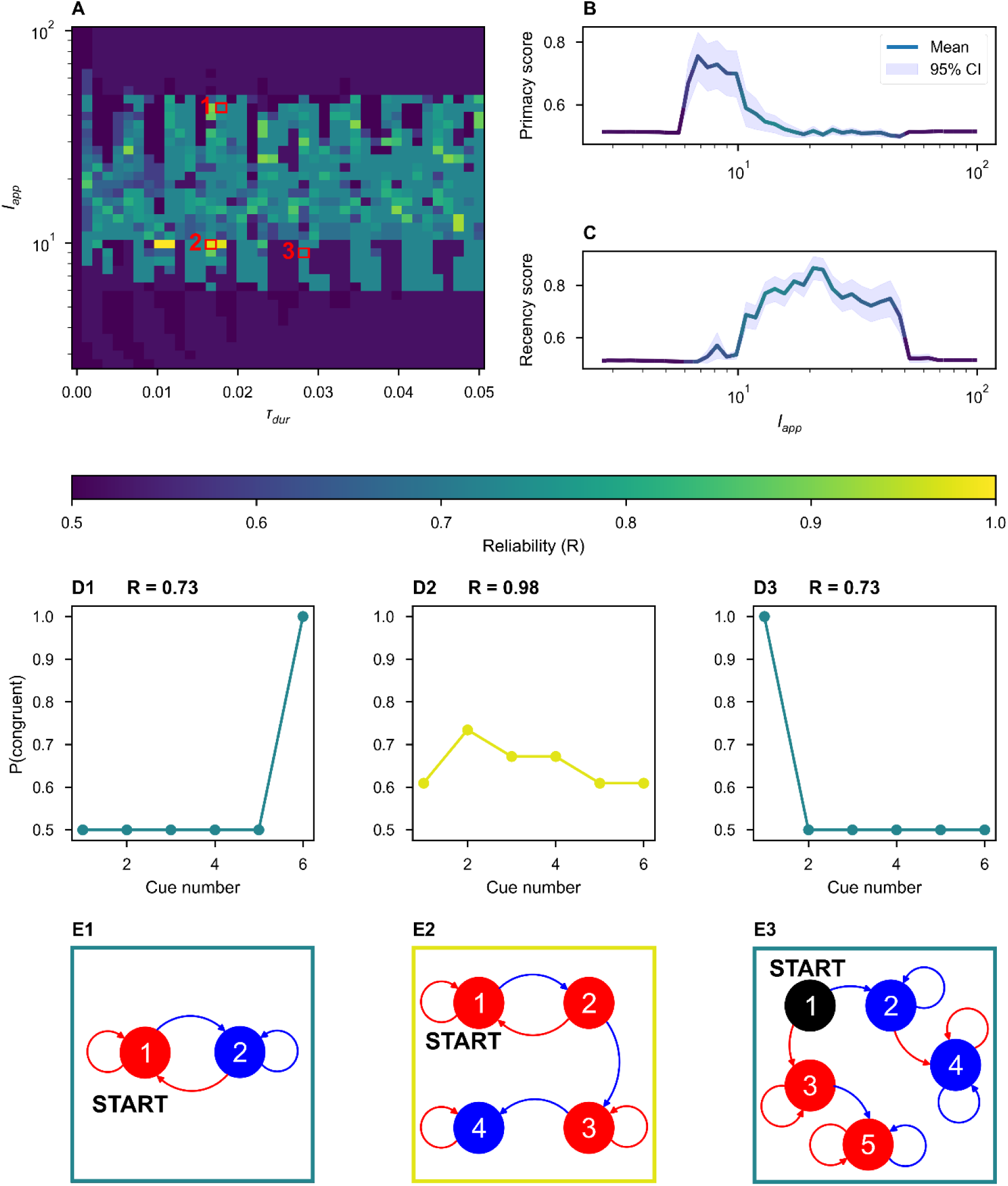
Sub-optimal state machines implemented by the ISN display primacy and recency. **(A)** The duration and amplitude of the applied stimulus were varied continuously (log axis for amplitude). The network reliability (Eq. 9) is displayed as the color. **(B-C**) The average primacy **(B)** and recency **(C**) scores as a function of the stimulus amplitude. The color of the line encodes the reliability of the network. Note that there is no clear relationship between reliability and amplitude. Weaker stimuli elicit primacy **(B)**, while stronger stimuli elicit recency **(C)**. **(D)** Examples of the computed congruence between cues at each position and the final network decision. **(D1)** A network implements a state machine that display recency. **(D2)** A high reliability network displaying minimal primacy and recency. **(D3)** A network displaying primacy. **(E)** State-machine diagrams corresponding to the examples in **(D)**. Node color encodes the direction of the decision for all sequences terminating in that state (blue: left, red: right, black: none).

We varied the amplitude and duration of the stimulus presented to the example network from Figure 5 and recorded the network’s reliability on the task (Fig. 6A). While the network and the stimulus weight vectors are fixed, changing the physical and temporal characteristics of the stimuli results in the implementation of new state machines that address a slightly different task (*i.e.,* the left-right task with a slightly different input alphabet). For a wide range of stimulus parameters, we found performance at baseline or above, though near perfect performance for only a small subset of stimuli.

Networks achieving below-perfect accuracy can display primacy or recency effects, for which we defined two continuous measures: the probability of congruence between the network’s final decision and first or last cue, respectively. Primacy arose, on average, when the stimulus is weak (Fig. 6B), while recency occurred, on average, when the stimulus is relatively strong (Fig. 6C). We observed a range of stimulus amplitudes for which any sequence discrimination was possible; outside of this range, network decisions were no better than chance, so no primacy nor recency were observed. Intuitively, when the stimulus is weak relative to internal network dynamics, a network is likely to get stuck in one of the first basins of attraction it enters, whereas when the stimulus is relatively strong, the final stimulus is likely to uniquely determine the final state, independently of the penultimate state such that history-dependent information is lost. We found no general relationship between the stimulus amplitude and state-machine reliability, most clearly (not) visible in Fig. 6B and 6C, where the color of the line corresponds to the average reliability over each amplitude. The perfect state machine exhibits neither primacy nor recency, so we might expect the highest performance at an intermediate amplitude, but no clear trends emerged to predict the performance from amplitude alone.

## 6 Heterogeneity in cross-connectivity

We were interested in whether heterogeneity in the cross-connections, while reducing the total number of attractor states from the 2^𝑁^ present in an appropriate homogeneous network, could at an optimal level, enhance robust network itinerancy. To test this, we varied the coefficient of variation in the cross-connectivity matrices, keeping the mean input to each unit constant. We drew weights from a gamma-distribution to more easily enforce Dale’s Law on large, random matrices. For each of these distinct networks, we tested a wide range of stimuli, as in Figure 4C. Figures 7B and C summarize network itinerancy across these stimuli for each degree of heterogeneity in the cross-connectivity.

**Figure 7.**
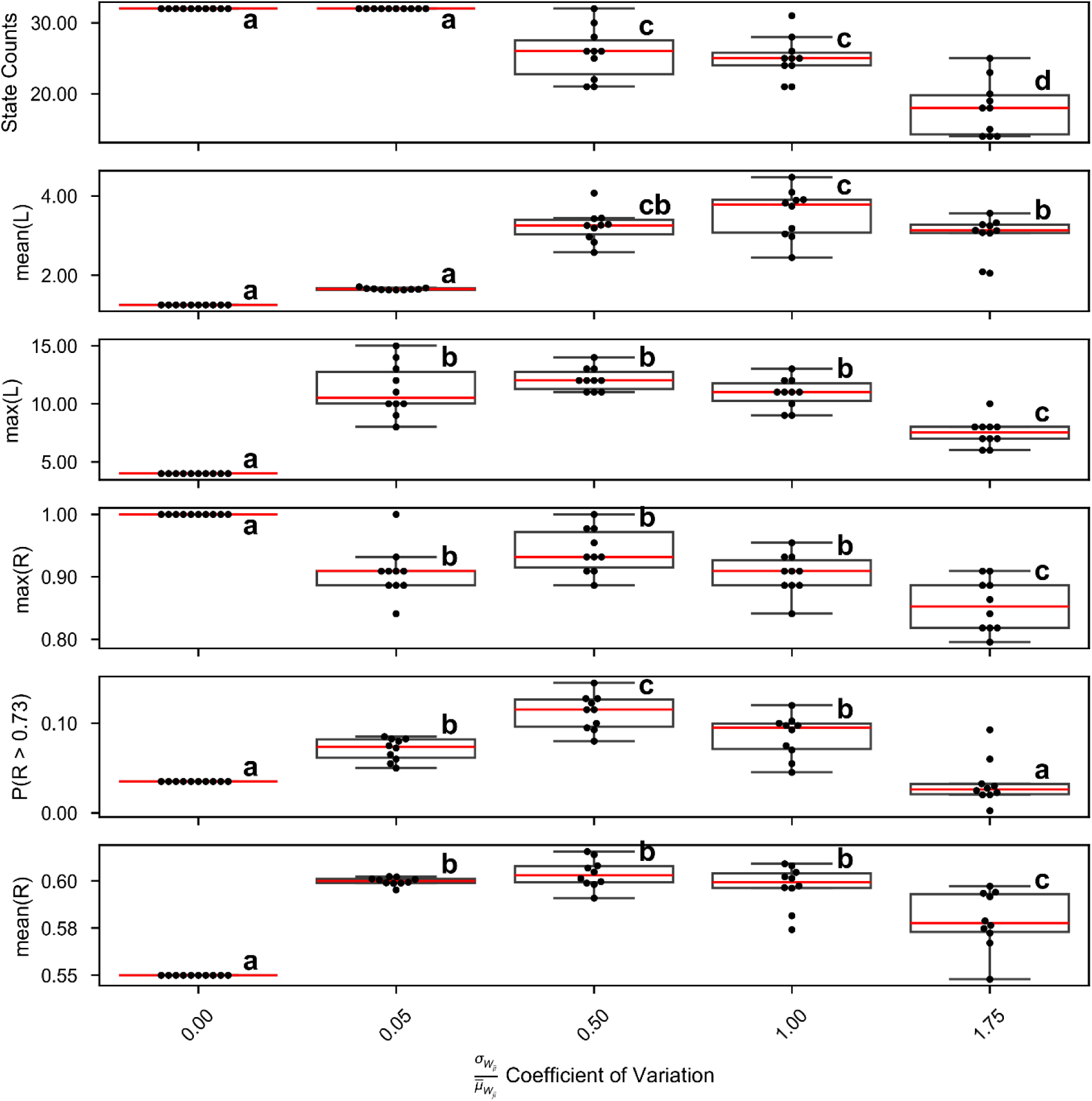
Heterogeneity enhances itinerancy and state-machine performance. The coefficient of variation of the random cross-connectivity is varied on the y-axis. For each level of variation, 10 unique connectivity matrices are generated with the correct statistics. **(A)** For each matrix, the number of attractor states is recorded. Increased heterogeneity decreases the number of attractor states observed. **(B-C)** For each matrix, which describe unique networks, a wide range of stimuli are applied and the length (L) of the longest itinerant path for each stimulus is recorded. **(B)** The mean longest path across all stimuli. **(C)** The global maximum path across all stimuli. **(D-E)** For each network, a wide range of stimuli are applied and the reliability on the left-right sequence discrimination task is recorded. **(D)** The highest reliability across all stimuli. **(E)** The percent of stimuli for which the network achieves above basline reliability. **(F)** The mean reliability across all tested stimuli. For all plots, compact letter display notation indicates statistically significantly different groups (ANOVA, Tukey HSD, 𝜶 = 𝟎. 𝟎𝟎𝟓 after Bonferroni correction).

For low variation, limited itinerancy is possible. Indeed, it can be logically demonstrated that a homogeneously connected network of identical rate units can produce a maximum itinerant walk of four transitions in response to a homogeneous input signal. The reasoning is as follows: suppose a network is in an attractor state with *n* pairs in the UP state (where 0 < *n* < *N*). Each active pair receives the same inputs from any stimulus and from internal connectivity, thus if a stimulus is sufficient to silence any one pair, it is sufficient to silence every pair in the UP state. The converse is true for quiescent pairs: any stimulus sufficient to make a silent pair active will flip every silent pair into the UP state. Thus, three possible stimulus-evoked state transitions are possible from our initial state: 1) all units flip states, resulting in a state with *N* – *n* pairs in the UP state, 2) all units transition to the UP state, or 3) all units transition to the DOWN state. Suppose the first transition occurs. From this new state, the response to the next stimulus is similarly limited. If the network transitions to an “all-UP” or “all-DOWN” state, it is, at best, limited to flipping back and forth between “all-UP” and “all-DOWN”. Thus, the longest possible itinerant path is from state *n*-UP, to (N-n)-UP, to all-UP/DOWN, to all-DOWN/UP, and finally to all-UP/DOWN again, a total of four transitions.

For excessively high variation, the network becomes more sparsely connected by high-weight cross-connections to maintain the same mean weight, and the number of possible states is reduced (Fig. 7A), which in turn reduces the maximum itinerancy (Fig. 7B-C).

We hypothesized that high-magnitude and robust itinerancy is valuable for network performance, but a direct, causal relationship is unproven. To investigate the correspondence between itinerancy and network performance, we again vary the heterogeneity of the random, weak cross-connections between pairs, as before. At each level of heterogeneity, we test network performance on a range of stimulus amplitudes and durations, keeping the optimal stimulus weights fixed. Again, network performance depends on a balance between sparseness and homogeneity. In a network with zero heterogeneity, the longest possible itinerant path is just four states. While this is technically sufficient to solve the task (cf Fig. 5C), it is exceedingly unlikely for a given mapping of the stimuli onto the network to elicit the correct responses. Thus, in these networks, the maximum reliability across all stimuli is a perfect 1, but the average reliability is low, indicating limited robustness to stimulus variability. In networks with excessive heterogeneity, the limited itinerancy and low state count similarly makes a reliable solution unlikely.

In summary, an optimal level of heterogeneity in the network can maximize itinerancy and maximize task performance. Too little heterogeneity leads to subsets of units responding identically to stimuli, so the system does not fully explore the vast repertoire of states, whereas increasing heterogeneity causes fixed points to collide, diminishing the total number of attractor states and reducing task performance.

## 7 Discussion

State-dependent responses to stimulation are necessary for many computational tasks.

Here, we demonstrated how inhibition-stabilization, in addition to stabilizing network activity at arbitrarily low values, can support dynamics that enhance state-dependent computation.

Specifically, inhibition-stabilization can produce transient oscillations around attractor states, which is sufficient for a repeated excitatory signal to both activate and silence neural populations, promoting robust itinerancy. From prior studies, we would expect that many forms of negative feedback can be tuned to produce oscillations sufficient for history dependence [11–13]. Nonetheless, inhibition-stabilized dynamics offer a simple, robust, and experimentally observed mechanism by which attractor states can be destabilized by excitatory input.

We enforce two constraints on an otherwise randomly connected attractor network composed of excitatory and inhibitory rate units: 1) strong connectivity within E-I rate unit pairs to produce inhibitory-stabilization and 2) homeostatic normalization of input weights to each rate-unit. Together with a moderate level of heterogeneity in cross-connectivity, these constraints are sufficient to produce a rich repertoire of structured dynamics that can be exploited to support state-machine-like computations. We demonstrated behaviorally realistic performance on a two-alternative forced-choice evidence accumulation task, including primacy and recency effects that depend on the magnitude of the stimuli.

### 7.1 Comparison to prior models

The models most similar to the ISN are the excitation-dominated attractor networks considered in [11–13, 29]. The excitation-dominated networks were composed of units with strong self-excitation, with either random weaker excitatory connections between units combined with a single global inhibitory unit connected all-to-all, or random excitatory/inhibitory connections between units with mean strength of zero (see also [30]). Qualitatively, the behavior of our ISN and these prior models are very similar: both possess a multiplicity of high-dimensional quasi-stable attractor states and network activity moves between these attractors in response to external stimulation. However, the differences in our ISN’s architecture result in crucial differences. First, by construction, inhibitory stabilization lowers the stable firing rates of the network to realistic, lower values. Second, we can achieve history-dependent responses without the assumption of strong, ubiquitous short-term synaptic depression, which was necessary for history dependence in these prior architectures.

A history-dependent response to incoming stimuli is essential for any task involving counting or integration of evidence. Indeed, we demonstrate the computational ability of the circuit by showing how it can account for the decision making and underlying neural activity in a task involving a numerical counting and comparison of two distinct stimuli [5, 11]. While transitions between discrete states are observed in such a counting task [5], whether they arise in more continuous integration-of-evidence tasks [6, 7, 31–34]—or whether a continuous model like the drift-diffusion model [35, 36] (or a combination of continuous diffusion then state transition) is more applicable—remains a matter of debate and ongoing study [37].

In their work in the field of vector-symbolic computing, Cotterret et al. [38] show that the weight matrix of a Hopfield network [39] can be constructed to produce pre-defined state-dependent transitions between arbitrary attractor states in response to inputs. In their model, each stimulus is represented by two extremely high-dimensional random input vectors that push the network between traditional Hopfield attractor states by traversing specially constructed intermediate “edge” attractor states, which encode the state-machine outputs. This construction contrasts with our encoding of stimuli in a single input vector and our treatment of “accepting states” as the encoding of network outputs. By their method, a sufficiently large network can be constructed to implement any arbitrary state machine, which is a significant advantage over the randomly constructed networks we considered here. An investigation of how realistic synaptic plasticity rules could convert the random attractor networks we analyze into task-specific state machines should be a fruitful avenue for further investigation.

Chandra et al. [40] use a fixed recurrent attractor network representing clusters of similarly-tuned neocortical grid cells as a “scaffold” or pool of available memory states, where the numerosity of attractor states scales with the number of grid modules. In their model of associative and episodic memory, a hippocampal-entorhinal circuit learns through Hebbian plasticity to map sensory inputs onto these attractor states, allowing the network to reconstruct noisy inputs and perform a spatial inference task without the catastrophic forgetting demonstrated by other Hopfield networks. While the spatial inference task demonstrates their network’s capacity for history-dependent responses, the mechanism for integration of a velocity vector over time is specific to grid cells. Clearly, non-spatial history-dependent behavior is possible, and it is not obvious how the demonstrated mechanism for history dependence might generalize to arbitrary computational problems.

Like Cotteret et al., our model implements arbitrary state machines, but, like Chandra et al., model dynamics are more strongly constrained by biology: network activity matches inhibition-stabilized dynamics, evolves in continuous time, and doesn’t rely on extremely high-dimensional attractor states for computation. The gap that remains to be closed is the implementation of an arbitrary, pre-specified state-machine by a combination of unsupervised and reinforcement learning in a biologically constrained attractor network: achieving the same capabilities as Cotteret et al. with higher fidelity to biology.

### 7.2 Cortical finite-state machines

In Section 5, we used the formal framework of finite-state automata to understand how transitions between persistent patterns of neural activity constitute primitive computations: as the network receives stimuli, it moves through attractor states in a manner that encodes information about the entire history of stimulation, ultimately settling in a final attractor state that can be read out as a decision. We show how a state transition function can be implemented purely by the dynamics of a randomly connected recurrent model of cortex. Cortex is crucial to cognitive processing, raising the question of whether the finite-state machine is the primary computational mechanism underlying cognition.

First, it must be observed that the cognitive abilities of human and some non-human animals appear to exceed the computational abilities of finite-state automata. FSMs are quite powerful: all digital computers, despite being theoretically modeled as Turing machines, can also be understood as FSMs and thus FSMs are sufficient for many tasks of practical interest [26]. On the other hand, because they have finite memory, many problems are not solvable by an FSM. FSMs definitionally cannot recognize context-free grammars, where valid sequences are constructed through recursive rules. For example, it is not possible to devise an FSM that only accepts palindromic sequences [26], nor one that can recognize valid sequences in a language composed of open and closed brackets, ex. “{[()[]]}”. Context-free grammars also occur in natural languages, such as with center-embedded sentences, ex. “The rat the cat the man bought chased ran.” [41]. Nonetheless, it has been demonstrated that humans [41], European starlings [42], and crows [43] can learn to recognize these kinds of context-free grammatical rules.

This evidence would seem to severely limit the explanatory power of the finite-state machine model of cognitive processing in cortex. However, it is worth noting that no group is perfect at performing these tasks [40–42], thus even if a state-machine solution is behaviorally sub-optimal, it may still be an accurate characterization of how cortex performs many computations in practice. Additionally, as might be expected, there are known limits on the length of context-free sequences that can be recognized by the organisms in each of these experiments. Humans, across seven languages, nearly entirely avoid doubly center-embedded structure when speaking, and in writing never use more than three nested center embeddings (as in the example above) [41]. Starlings [42] and crows [43] display decreased accuracy in discriminating between sequences as the sequence length increases.

These observations address two larger points. First, while the finite memory of an FSM limits the scope of languages it can recognize in theory, in practice there are no cognitive tasks that require infinite memory and certainly no such tasks that any organism can perform. Second, relatedly, an FSM can approximate the behavior of a more powerful model like a pushdown automaton if bounds are imposed on the language. A pushdown automaton is an FSM with access to an infinite stack, which allows the automaton to recognize a wider class of languages, including center-embedded syntaxes [26]. If we impose practical constraints on the problem by bounding the length of the strings, an FSM can solve the task simply by representing every possible sequence. Conversely, if we impose constraints on the pushdown automaton by bounding the size of the stack, the automaton will only recognize sequences up to a finite length. Notably, in the latter scenario, the pushdown automaton is equivalent to a finite-state machine because the number of possible stack configurations is finite. Thus, an FSM can at least approximate a solution to any real-world cognitive task, which aligns with the documented limitations of human and non-human animal abilities to recognize context-free grammars.

The real challenge for the FSM as a model of cortical computation is the question of how an attractor network can learn to implement a state machine via a mixture of unsupervised and reinforcement learning. That is, given a subset of all possible stimulus sequences, how does a network learn a history-dependent encoding that generalizes the decision rule to unseen stimulus sequences? There are at least three places where synaptic plasticity could act on an attractor network to tune the computation being performed.

First, the weights of the recurrent attractor network could themselves be adjusted, though we would expect that the weights within inhibition-stabilized subnetworks would remain relatively constant to maintain the benefits of inhibitory-stabilization. Here, changes in network parameters shift the geometry of the attractor wells, resulting in changes in the state-transition function.

Second, the weights of the inputs to the network could be adjusted. We showed that random networks can, just by chance, display dynamics that separate sequences based on the frequencies of their (random) component stimuli. Modifying the parameters of the input to the attractor network could take advantage of the rich dynamical repertoire of a random attractor network to change the state-transition function and perform an entirely new computation. This approach can be thought of as a history-dependent adaptation of Chandra et al.’s [40] mechanism for the association of arbitrary inputs with fixed, random attractor states. In this view, when a network shifts to a new task, only the input to the neocortical network need change, allowing the same attractor network to implement multiple computations without changing the 𝑂(𝑁^2^) recurrent weights.

Finally, the output function that “reads out” the decision from the final attractor network state could adjust to reinterpret the sequence encodings from the cortical network to enforce a different decision rule. For instance, if we assume a network that perfectly solves the task considered in Section 5, it is also possible to learn the reverse task, “turn towards the direction with the fewest cues,” just by changing the output function. Indeed, given a fixed attractor network with *k* reachable states, the number of possible non-isomorphic output functions (representing unique sequence discrimination tasks), is given by the *k*-th Bell number: the number of ways to partition the *k* reachable states.

Likely, some combination of these changes is necessary, but it remains to be demonstrated that a biologically plausible learning rule could allow a cortical attractor network to learn an arbitrary FSM.

### 7.3 Experimental predictions

Transitions between high-dimensional persistent activity states has already been observed in cortex during decision-making behavior [5, 7, 34]. Our work here makes additional testable predictions about how these transitions might be occurring.

First, our hypothesis is that transient oscillations are what permit transitions between more active and less active attractor states in response to an excitatory stimulus. We would expect to see high amplitude transient oscillations in activity during state transitions, which is measurable with intercellular recordings.

Another prediction is that the salience of a stimulus might induce primacy or recency effects. If a stimulus elicits a stronger neural response in the primary sensory region corresponding to the stimulus, we would predict a recency effect. Conversely, a weak neural response in primary sensory cortex would predict a primacy effect.

Additionally, while not studied here, the effects of internally- or externally-generated noise could be studied by treating the attractor network as an implementation of a non-deterministic finite state machine, where the state-transition function outputs a probability distribution over states [26]. Each sequence of inputs would therefore be mapped by a network onto a distribution of final attractor states, over which our reliability measure could be generalized to an “expected reliability.” We would expect that noise would have the most deleterious impact on network performance during state transitions: while the network is in an attractor state, small perturbations will decay over time, but between attractor states, noise could result in erroneous state transitions. We would predict that noisy perturbations at the time of stimulus presentation would be more impactful than noisy perturbations between stimuli.

In summary, we have shown that an inhibition-stabilized attractor network can implement history-dependent information processing, which depends on transient oscillations facilitating transitions between attractor states, implying that primacy/recency effects depend on stimulus salience and that cortical networks are most sensitive to perturbation during stimulus presentation.

## Acknowledgements

The authors are grateful to Ann Kennedy for her comments on the manuscript. This work was supported by funding from NIH-NINDS via research grant R01NS104818 to PM, from NIH-NIDA via training fellowship R90DA060341 to CH, the Blavatnik Family Foundation and Goldwater Scholarship Foundation for scholarships to CH.

