## Supplemental Materials for "Inhibitory-stabilization is sufficient for history-dependent computation in a randomly connected attractor network"

### A. Dynamics of a single pair

The dynamics of a single inhibitory-stabilized pair without external input are described by the system of differential equations:

$$\tau \dot{r}_E = g_E(-r_E + W_{EE} \cdot r_E + W_{IE} \cdot r_I - \theta_E)$$

$$\tau \dot{r}_I = g_I(-r_I + W_{EI} \cdot r_E + W_{II} \cdot r_I - \theta_I)$$

$$g_{E/I}(x) = \begin{cases} x, & r_{E/I} > 0 \\ max(x, 0), & r_{E/I} = 0 \end{cases} \quad (1)$$

All parameters are the same as defined in the methods section and we assume Dale's law is enforced on the weight parameters. Here, we more explicitly denote the enforcement of "sticky" boundary conditions with the function *g*. The $r_E$- and $r_I$- nullclines, $\mathbf{N_E}$ and $\mathbf{N_I}$ **(Fig. 1)**, are given by:

$$\boldsymbol{N_E} = \{(r_E, r_I) \in \mathbb{R} \mid (r_E > 0\ AND\ r_I = \frac{(W_{EE} - 1)r_E - \theta_E}{-W_{IE}})\ OR\ (r_E = 0\ AND\ r_I \geq \frac{\theta_E}{W_{IE}})\}$$

$$\boldsymbol{N_I} = \{(r_E, r_I) \in \mathbb{R} \mid (r_I > 0\ AND\ r_I = \frac{W_{EI} r_E - \theta_I}{1 - W_{II}})\ \text{OR}\ (r_I = 0\ AND\ r_E \leq \frac{\theta_I}{W_{EI}})\}\ \#(2)$$

In each line, the second condition after the "OR" describe the threshold nonlinearity: line-segments along the $r_I$- and $r_E$-axes we will call the "boundary lines". The origin is a fixed point only when (0,0) is an element of both nullclines, which can only occur when the boundary lines include the origin (Fig 1B,D). Explicitly:

$$0 \geq \frac{\theta_E}{W_{IE}}\ AND\ 0 \leq \frac{\theta_I}{W_{EI}} \quad (3)$$

which simplifies to $\theta_E \geq 0$ and $\theta_I \geq 0$ (multiplying by $W_{IE}$ flips the inequality). When the threshold values are strictly positive, the origin is locally asymptotically stable: for sufficiently small $r_E$ and $r_I$, equation (1) is dominated by $\theta_E$ and $\theta_I$, which because they are positive, push the rates to zero, where they stick:

$$\tau\dot{r}_E = g_E(-\theta_E) = 0$$

$$\tau\dot{r}_I = g_I(-\theta_I) = 0 \qquad (4)$$

The UP fixed point is the intersection of the "interior" nullcline segments – those defined by the first conditions in equation (2) (Fig 1A). Its coordinates are given by:

$${r_E}^* = -\frac{W_{IE}(\theta_I) + (1 - W_{II})(\theta_E)}{d}$$

$${r_I}^* = -\frac{W_{EI}(\theta_E) + (1 - W_{EE})(\theta_I)}{d}$$

$$d = (W_{EE} - 1)(W_{II} - 1) - W_{EI} \cdot W_{IE} \qquad (5)$$

The Jacobian of equation (1), evaluated at the non-zero fixed point is:

$$\frac{1}{\tau} \cdot \begin{bmatrix} W_{EE} - 1 & W_{IE} \\ W_{EI} & W_{II} - 1 \end{bmatrix}. \qquad (6)$$

Thus, the trace and determinant of the Jacobian are:

$$Tr = \frac{1}{\tau}(W_{EE} + W_{II} - 2)$$

$$\Delta = \frac{d}{\tau^2} \qquad (7)$$

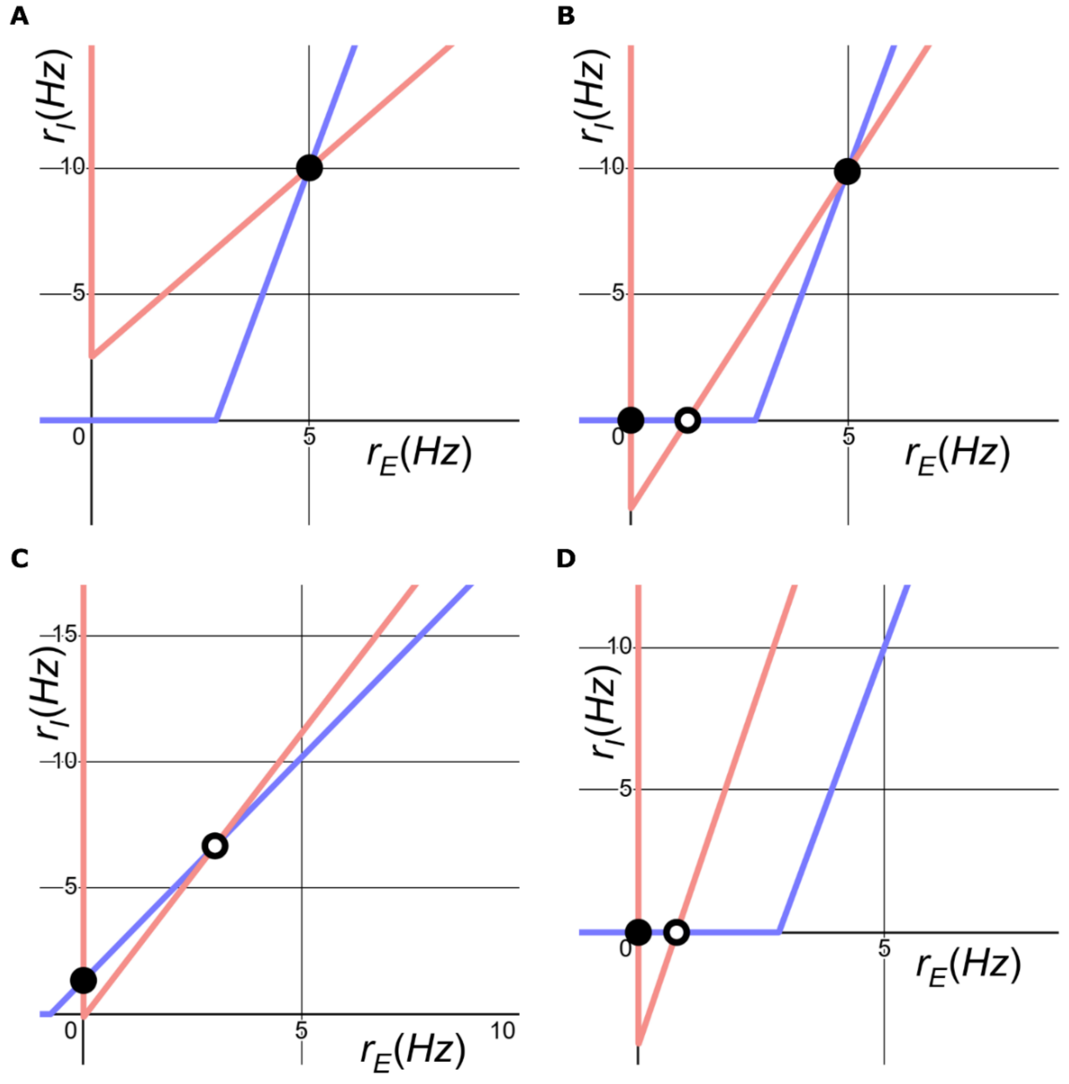


**Supplementary Figure 1: Examples of four of the possible dynamical phases of an E-I pair.** **A.** Inhibition-stabilized. **B.** Inhibition-stabilized and bistable. **C-D.** Stable state with E-unit off, active state unstable, not inhibition-stabilized. Parameters used are as follows: **A.** $\theta_E = -2$, $\theta_I = 20$, $W_{EE} = 2$, $W_{EI} = 7$, $W_{IE} = -0.7$, $W_{II} = -0.5$; **B.** $\theta_E = 2.5$, $\theta_I = 20$, $W_{EE} = 2.9$, $W_{EI} = 7$, $W_{IE} = -0.7$, $W_{II} = -0.5$; **C.** $\theta_E = 0$, $\theta_I = -4$, $W_{EE} = 2$, $W_{EI} = 7$, $W_{IE} = -0.7$, $W_{II} = -4.5$; **D.** $\theta_E = 2.5$, $\theta_I = 20$, $W_{EE} = 43.5$, $W_{EI} = 7$, $W_{IE} = -0.66$, $W_{II} = -0.5$.

The conditions for the stability of the UP fixed point are therefore, from the trace and determinant respectively:

$$W_{EE} + W_{II} < 2$$

$$d = (W_{EE} - 1)(W_{II} - 1) - W_{EI} \cdot W_{IE} > 0 \tag{8}$$

Note that the condition derived from the determinant can be understood as a relationship between the slopes of the interior segments of the nullclines:

$$\frac{W_{EE} - 1}{W_{IE}} < \frac{W_{EI}}{1 - W_{II}} \tag{9}$$

That is, one condition for stability is that the gain of the inhibitory nullcline be greater than the gain of the excitatory nullcline. The set of conditions for existence and stability of the two fixed points described above are necessary conditions for the bistability we study in the paper. Supposing that all the above conditions are met, we can derive a fourth condition that guarantees that there are two stable fixed points: one at the origin and an UP-state $(r_E{}^*, r_I{}^*) \in (0, \infty)^2$.

First, consider the fixed point:

$$r_E{}^* = -\frac{W_{IE}(\theta_I) + (1 - W_{II})(\theta_E)}{d} > 0$$

$$r_I{}^* = -\frac{W_{EI}(\theta_E) + (1 - W_{EE})(\theta_I)}{d} > 0 \tag{10}$$

Because the determinant is positive by assumption, d is also positive. Thus, we can simplify:

$$(W_{II} - 1)\theta_E - W_{IE}\theta_I > 0$$

$$(W_{EE} - 1)\theta_I - W_{EI}\theta_E > 0 \tag{11}$$

Notice that ($W_{EE}$ – 1) > 0. Otherwise, the second inequality is impossible because all involved parameters are positive by the assumption of Dale's Law. Rearranging,

$$\theta_I > \frac{(W_{II} - 1)\theta_E}{W_{IE}}$$

$$\theta_I > \frac{W_{EI}\theta_E}{(W_{EE} - 1)} \tag{12}$$

And so,

$$\theta_I > max\{\frac{(W_{II} - 1)}{W_{IE}}, \frac{W_{EI}}{(W_{EE} - 1)}\} \theta_E$$

By Dale's law and equation (11), both possible coefficients for $\theta_E$ are positive, thus

$$\theta_I > \theta_E \tag{13}$$

Together, equations (8) and (13) are necessary and sufficient for the bistability studied in this paper (Supp. Fig. 1B).

There two more possible fixed points – those which results from the intersection of the "interior" and "boundary" segments of the nullclines (Supp. Fig. 1B-D). In the bistable, inhibitory-stabilized regime (Supp. Fig. 1B), one of these fixed points exists between two stable fixed points: the origin and the UP state, and is therefore an unstable node or a source (Supp. Fig. 1B). In the cases where these fixed points are not ringed by attractors, the network is not inhibition-stabilized (Supp. Fig. 1C-D). Thus, explicit treatment of these fixed points is unnecessary for our inhibition-stabilized systems.

#### B. Fiducial Parameter sets

The "fiducial" parameter sets for each figure are contained in the following table. A dash indicates the parameter is not relevant to the figure. For the final four rows, the overbar indicates the mean of the cross-connections of that type and sigma indicates the standard deviation of those cross-connections. The exact values of the cross-connections can be found in the GitHub repository, `model` folder.

| | Figures 1-2 | Figure 3 | Figures 4-7 |
|---|---|---|---|
| $\tau$ | $10\ ms$ | $10\ ms$ | $10\ ms$ |
| $W_{EE}$ | 16.47 | 16.47 | 11.979 |
| $W_{EI}$ | 51.60 | 51.60 | 43.644 |
| $W_{IE}$ | −7.20 | −7.20 | - 4.751 |
| $W_{II}$ | −16.55 | −16.55 | - 10.343 |
| $\theta_E$ | 5.34 | 5.34 | 7.390 |
| $\theta_I$ | 82.43 | 82.43 | 104.790 |
| $\overline{W_{EE}}$ | - | $0.0107 * 5/N$ | 0.0107 |
| $\overline{W_{EI}}$ | - | $0.00327 * 5/N$ | 0.00327 |
| $\overline{W_{IE}}$ | - | $-0.283 * 5/N$ | −0.283 |
| $\overline{W_{II}}$ | - | $-0.00140 * 5/N$ | −0.00140 |
| Coefficient of variation | - | 0.5 | 0.5 |

**Supplementary Table 1: Fiducial parameter sets for each figure.** Parameters with overlines and the coefficient of variation correspond to cross-connections between pairs.

84 **C. Examples for Fig 2 history-dependence**

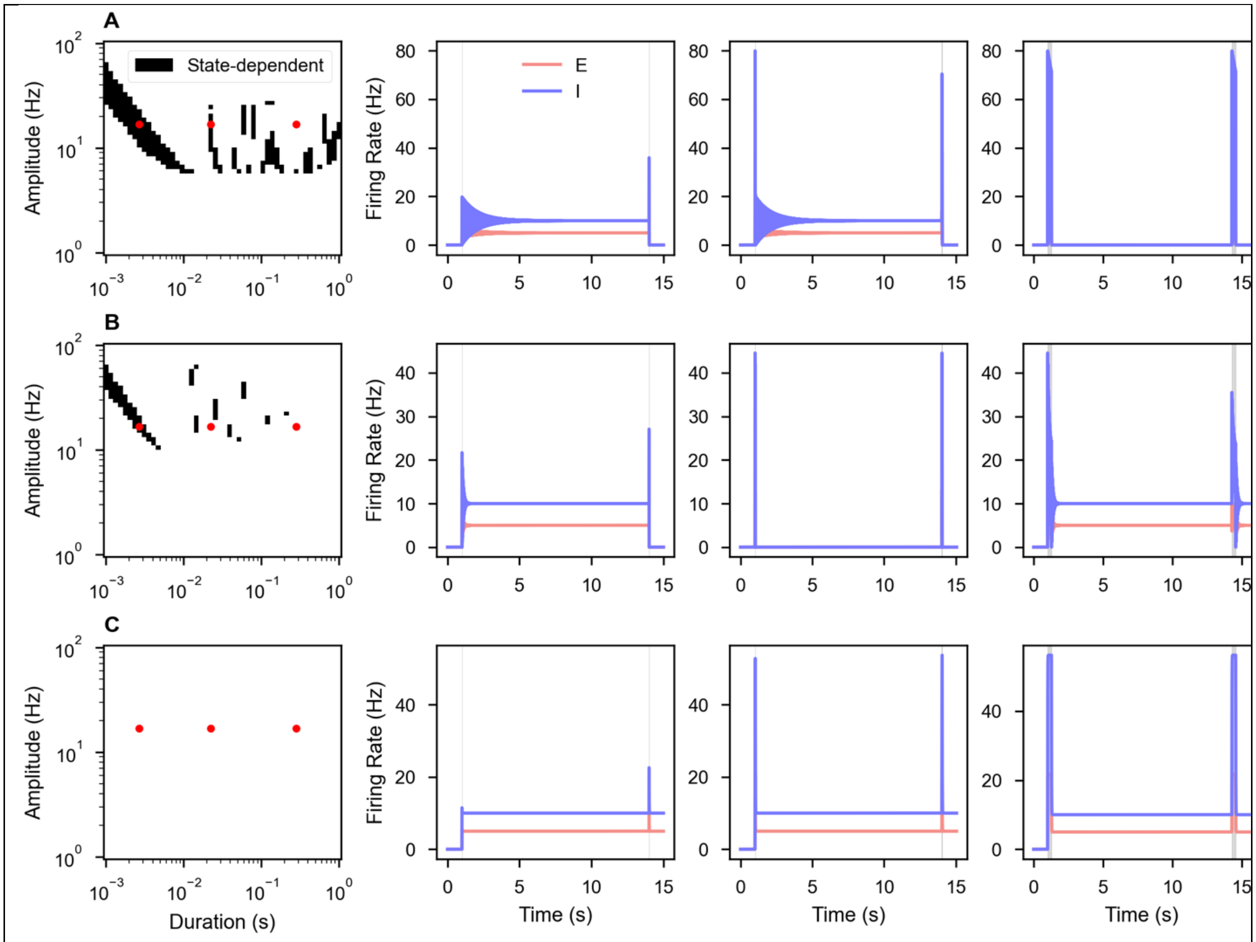


**Supplemental Figure 2. Dynamics of network in response to stimuli of different durations.** Rows A-C correspond to the networks in panels A-C of Figure 2, with network parameters depicted and labeled in Figure 2D. Each successive column corresponds to stimuli of increasing duration, indicated by a red solid circle in the first column. A repeated stimulus successively switches the network from quiescent to active and then from active to quiescent in **A)** the first two examples, in **B)** just the first example, and in **C)** not at all. Note that failures can arise by a stimulus being unable to produce an active state (row **A**, strongest stimulus and row **B**, middle stimulus) or by being unable to produce quiescence from the active state (row **B**, stongest stimulus and row **C**, all stimuli.)

**D. Method for continuous variation of trace and determinant**

In Figures 2 and 3, we continuously varied the trace (Tr) and determinant (Δ) of the UP state in a bistable inhibition-stabilized pair, while holding the stable firing rates ($r_E$* and $r_I$*) at the UP-state constant (denoted: $r_E^{des}$ and $r_I^{des}$). To enforce bistability, we also fixed the firing thresholds of the rate units at $\theta_E^{des}$ and $\theta_I^{des}$ such that $\theta_I^{des} > \theta_E^{des} > 0$. The timescale, τ, is fixed at 10 ms, as for all simulations.

Thus, the problem is to find the connectivity weights $W_{EE}$, $W_{II}$, $W_{EI}$, and $W_{IE}$ such that:

$$Tr = \frac{1}{\tau}(W_{EE} + W_{II} - 2) = Tr^{des}$$

$$\Delta = \frac{d}{\tau^2} = \Delta^{des}$$

$$r_E{}^* = \frac{W_{IE}\theta_I + (1 - W_{II})\theta_E}{d} = r_E{}^{des}$$

$$r_I{}^* = \frac{W_{EI}\theta_E + (1 - W_{EE})\theta_I}{d} = r_E{}^{des}$$

$$d = (W_{EE} - 1)(W_{II} - 1) - W_{EI} \cdot W_{IE} \qquad (14)$$

We imposed the additional constraints that 1) excitatory connections have positive value and inhibitory connections have negative value and 2) connections be non-zero, which we enforce practically by requiring all weights be greater than $W_{min}$ = 0.001. We implemented these constraints by modifying the problem, defining four new independent variables, $\omega_{xx}$ where x ∈

{E, I} such that $W_{xx} = \omega_{xx}^2 + W_{min}$. We found that squaring $\omega_{xx}$ resulted in more reliable convergence than the absolute value. Note that equations (14)form a nonlinear system of four equations with four unknowns. Thus, it is appropriate to use a numerical root solver to find a solution to the system:

$$\mathbf{F}^{*}(\omega_{EE}, \omega_{II}, \omega_{EI}, \omega_{IE}) = \begin{bmatrix} Tr - Tr^{des} \\ \Delta - \Delta^{des} \\ r_E{}^{*} - r_E{}^{des} \\ r_I{}^{*} - r_I{}^{des} \end{bmatrix} = \mathbf{0} \qquad \mathbf{(15)}$$

Convergence is not guaranteed. To increase the odds of convergence, we initialize 1000 different random initial conditions for the root solver. We slowly increase the range of the random number generator over successive initializations, which we found resulted in more reliable convergence.

#### E. Logistic vs linear psychometric curves

We fit logistic and linear curves to the psychometric data in Figure 5 of the form:

$$y_i = ax_i + b + \varepsilon_i, \varepsilon \sim N(0, \sigma^2)$$

and

$$y_i = \frac{c}{1 + e^{-a(x_i - b)}} + \varepsilon_i, \varepsilon \sim N(0, \sigma^2), \qquad (17)$$

where a, b, and c parameterize the curves and σ parameterizes the distribution of residuals, assumed to be normal for the purposes of likelihood estimation. We fit the curves to all psychometric data for networks achieving greater than 0.73 reliability. We compute the sum of the squares of the residuals (RSS), Akaike Information Criteria (AIC), and Bayesian Information Criteria (BIC) for both models, reported in the Table 2 below:

| | RSS | AIC | BIC |
|---|---|---|---|
| Linear | 30.231 | -987.6 | -972.4 |
| Logistic | **26.383** | **-1147.6** | **-1127.3** |

**Supplementary Table 2. Comparison of logistic and linear regression on psychometric curves.** Rows: model. Columns: metrics. Bold number in each column indicates the best score.

The logistic model has lower RSS, indicating better goodness-of-fit. Additionally, by both information criteria, which penalize the additional degree of freedom afforded to the logistic regression, the logistic curve is a better model (lower is better). The logistic nature of the psychometric curves, as opposed to linear, is characteristic of history-dependent computation. Both curves are plotted against the raw data (Supplemental Fig 2):

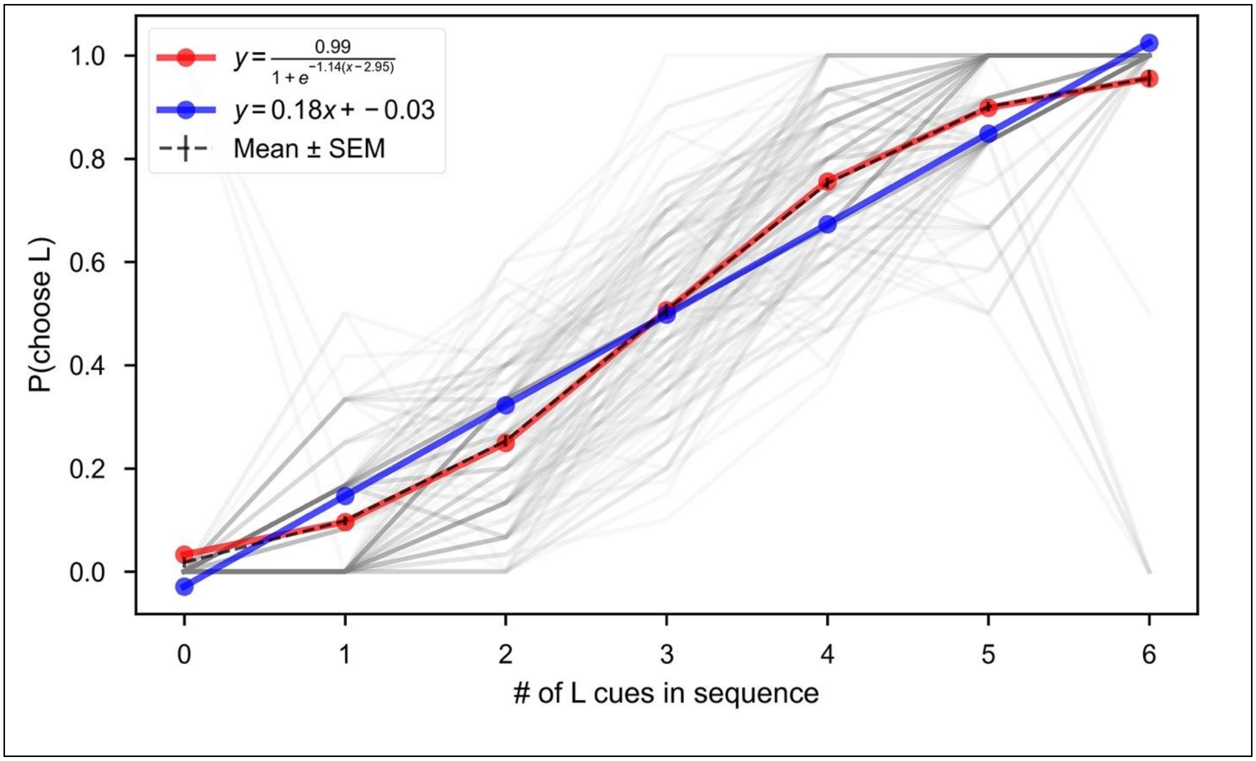

**Supplemental Figure 3. Logistic and linear fits to psychometric data.** Logistic fit (red) is significantly better than linear fit (blue) according to both AIC and BIC. Mean and SEM of the data are indicated by the black dashed lines, demonstrating the close agreement with the logistic fit.
